# Propagation of spike timing and firing rate in feedforward networks reconstituted *in vitro*

**DOI:** 10.1101/151134

**Authors:** Jérémie Barral, Xiao-Jing Wang, Alex Reyes

## Abstract

The manner in which information is transferred and transformed across brain regions is yet unclear. Theoretical analyses of idealized feedforward networks suggest that several conditions have to be satisfied in order for activity to propagate faithfully across layers. Verifying these concepts experimentally in networks has not been possible owing to the vast number of variables that must be controlled. Here, we culture cortical neurons in a chamber with sequentially connected compartments, optogenetically stimulate individual neurons in the first layer with high spatiotemporal resolution, and monitor the subthreshold and suprathreshold potentials in subsequent layers. In the first layer, a brief stimulus with different temporal precisions resulted in the modulation of the firing rate. This temporal to rate transformation was propagated to other layers as a sustained response, thereby preserving rate information. This novel mode of propagation occurred in the balanced excitatory-inhibitory regime and is mediated by NMDA-mediated synapses activated by recurrent activity.

---

Information is encoded in the nervous system as action potentials generated via the coordination of many networks in the different regions of the brain. Interactions between regions occur via feedforward architectures, which are embedded within the complex circuitry of the nervous system. These include, for example, sequential brainstem nuclei in sensory systems and the path from thalamus to the cortical layers. What signals are propagated through feedforward networks is still under debate. Information may either be represented as the average number of spikes per unit time (rate coding) (Newsome, Britten et al. 1989, Georgopoulos, Taira et al. 1993) or by their precise timing (temporal coding) (Riehle, Grun et al. 1997). The nervous system may also utilize a combination of both strategies and there appears to be a continuum between these two extreme coding schemes (Kumar, Rotter et al. 2010).

The firing rate can be modulated by features of the stimuli and thus contains information (Hubel and Wiesel 1962, Ahissar, Sosnik et al. 2000, Lu, Liang et al. 2001, Gao and Wehr 2015, Gao, Kostlan et al. 2016). In cortex, there are neurons that fire tonically with responses that outlast the duration of the sensory input in different modalities such as visual (Reinhold, Lien et al. 2015), auditory (Schroeder, Lindsley et al. 2001, Wallace and Palmer 2008), or somatosensory (Ahissar, Sosnik et al. 2000, Ahissar, Sosnik et al. 2001). This is perhaps most evident in cortical areas involved with working memory where a transient stimuli (cue) evokes firing responses (hold period) that are long lasting and in some cases persistent (Miller, Li et al. 1993, Brody, Hernandez et al. 2003). Modeling studies suggest that recurrent activity and/or NMDA-mediated synapses underlie the prolonged activity (Wang 1999). Moreover, in progressively higher order or downstream neurons, the firing patterns become more rate-like as transient components become less prominent (Ahissar, Sosnik et al. 2000, Lu, Liang et al. 2001, Gao and Wehr 2015).

The conditions for propagating rate signals across multiple layers are still poorly understood. The feedforward architecture places specific constraints on the signals that can propagate. Analyses of feedforward network consisting only of excitatory neurons with random connectivity suggested that only synchronous activity propagated. Activating a sufficiently large number of neurons in the first layer within a narrow temporal window (termed pulse packets) caused neuronal firing to become more synchronous in the subsequent layers whereas increasing temporal jitter and/or decreasing the number (broad, short packets) caused activity to rapidly dissipate (Aertsen, Diesmann et al. 1996, Diesmann, Gewaltig et al. 1999, Reyes 2003, Vogels and Abbott 2005, Kumar, Rotter et al. 2008). From a neural code perspective, the development of synchronous activity degrades rate signals (Zohary, Shadlen et al. 1994, Mazurek and Shadlen 2002, Litvak, Sompolinsky et al. 2003).

Asynchronous activity and hence rate information can propagate in feedforward networks under certain conditions. In simulations with sparse but strong synaptic coupling, networks exhibit less synchrony due to low shared connectivity between layers (van Rossum, Turrigiano et al. 2002, Vogels and Abbott 2005). However, complete asynchrony, a condition for rate propagation, was achieved either in the presence of noisy background inputs (van Rossum, Turrigiano et al. 2002, Litvak, Sompolinsky et al. 2003) or by embedding feedforward networks with strong connections within a recurrent network (Aviel, Mehring et al. 2003, Mehring, Hehl et al. 2003, Tetzlaff, Buschermohle et al. 2003, Vogels and Abbott 2005, Kumar, Rotter et al. 2008, Jahnke, Memmesheimer et al. 2013, Chenkov, Sprekeler et al. 2017). Modulating the relative timing or balance between excitation and inhibition provides a means for selectively propagating temporal or rate signals (Vogels and Abbott 2009, Kremkow, Aertsen et al. 2010, Shinozaki, Okada et al. 2010).

Here, we examined signal propagation in *in vitro* cultures of excitatory (E) and inhibitory (I) cortical neurons grown in a chamber with multiple compartments or layers arranged in a feedforward manner. Excitatory neurons in the first layer expressed channelrhodopsin and were individually stimulated with a specified spatio-temporal pattern while activity of neurons in subsequent layers was recorded. We find that the networks transformed temporal information into firing rate and can support and propagate sustained firing that far outlasts the stimulus. Firing rate levels are preserved in all layers. Analyses of subthreshold potentials suggest rate propagation occurs in the balanced E-I regime and is mediated by NMDA-mediated potentials enhanced by recurrent activity. Importantly, the time course of firing and the transformation across layers recapitulate those documented *in vivo*.

## Results

### Feedforward network *in vitro*

To examine experimentally the conditions for propagation of activity across layers, we cultured cortical neurons in chamber with interconnected compartments arranged in a feedforward manner (Fig. 1a,b). Neurons were cultured in 4 rectangular compartments (0.7 × 6 mm^2^) separated by 0.4 mm spacers, which were then removed after 24 hours to allow bidirectional growth of axons. At 14-21 days *in vitro* (DIV), neurons grew primarily in the compartments (henceforth termed layers) and formed recurrent and bidirectional synaptic connections within and across layers (see Supplementary Fig. 1). Previous studies showed that intrinsic and synaptic properties and the relative proportion and synaptic connection architecture between excitatory (E) and inhibitory (I) cells were similar to those measured *in vitro* (Barral and Reyes 2016, Nikitin, Bal et al. 2017).

**Figure 1:**
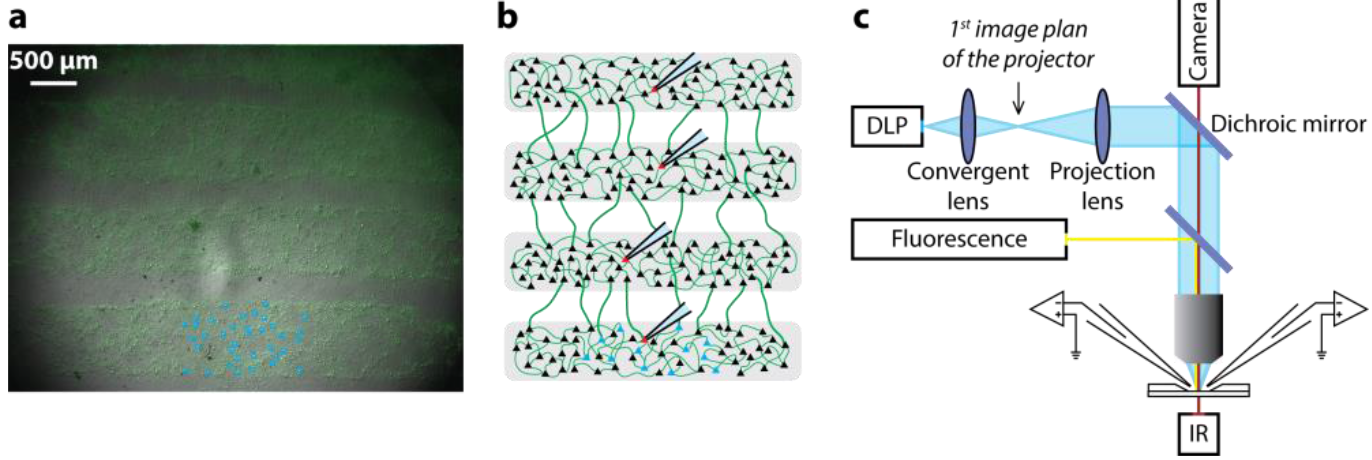
Building and stimulation of a feedforward network *in vitro*. **a.** Networks were visualized with IR-DIC and fluorescent microscopy. **b.** Schematic of the feedforward network. Spikes were recorded from 4 neurons in sequential layers. **c.** Using a DLP projector mounted on a microscope, brief light pulses (blue boxes in **a**) were delivered to neurons (in layer 1) expressing ChR2 and a fluorescent tag (green).

To characterize the connectivity patterns between neurons within and across layers, we performed paired whole-cell recordings and estimated the connection probabilities between cells. The connection probability between neurons within a layer (*P*_*c*_ = 0.3 ± 0.06; mean ± SEM; n = 50, tested connections; 533 ± 132 μm apart; mean ± SD) was comparable to those between neurons in two adjacent layers (*P*_*c*_ = 0.23 ± 0.04, mean ± SEM; n = 100 tested connections; 666 ± 140 μm apart; mean ± SD). The connection probabilities appeared to be determined only by the distance between neurons and were readily predicted from the connection probability profiles of neurons grown in a single compartment (see Supplementary Fig. 1 and ref. (Barral and Reyes 2016)). Given the long distances between chambers (~1.1 mm), connections across non-adjacent layers were rare so that propagation occurs sequentially and did not ‘skip’ layers (see simulations in Supplementary Fig. 2a). Both excitatory (E) and inhibitory (I) connections occurred across layers; hence, the architecture resembled more that of neurons across cortical layers than between neurons in different brain regions where inhibition is local.

### Propagation of pulse packets in a feedforward network

To examine propagation of activity across layers, we expressed channelrhodopsin (ChR2) in excitatory neurons. We then optically stimulated neurons in the first layer using a computer-controlled Digital Light Processing (DLP) projector to deliver independent blue light pulses (Fig. 1c; see (Barral and Reyes 2016, Barral and Reyes 2017)) as follows. Approximately 20-40 neurons within a 0.7 × 1.5 mm^2^ region were marked for stimulation with regions of interests (ROIs, Fig. 1a). A single brief blue light pulse (5 ms) was delivered to each neuron to evoke action potentials. Here, pulses were synchronized (Fig. 2) and spikes and membrane potential of neurons in layer 1-4 were recorded using cell-attached and whole-cell recordings, respectively.

**Figure 2:**
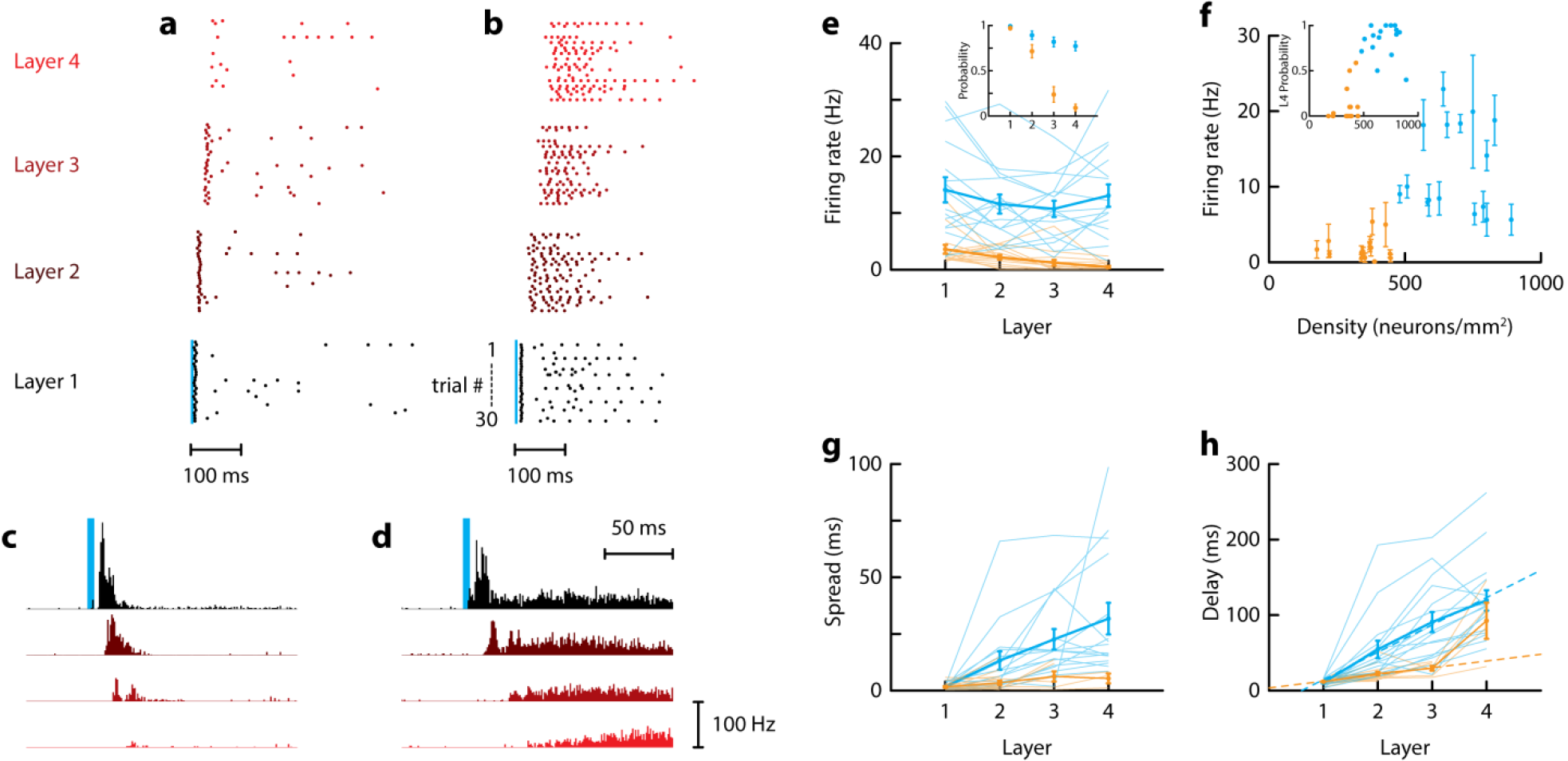
Activity propagation in feedforward networks reconstituted *in vitro*. **a.-b.** Dot rasters showing spikes recorded from a neuron in each layer in cell attached mode in response to synchronized activation of neurons in the first layer (30 repetitions of the light stimuli). Results shown for a sparse (**a**) and a dense (**b**) network. **c.-d.** Average peristimulus time histogram (PSTH) of neurons in sparse (**c**) or dense (**d**) networks. In each network, a single neuron was recorded in each layer (from black to red for layer 1 to 4, respectively; same color code in **a-b**). **e.** Firing rate in the 500 ms after the stimulus vs recorded layer in sparse (orange) and dense (blue) networks. Inset: Spike probability vs recorded layer (the spike probability was defined as the probability to evoke at least 1 spike in the 500 ms following the stimulation). **f.** Firing rate averaged over the 4 layers as a function of network density. Inset: spike probability in the 4^th^ layer as a function of density. **g.-h.** Temporal spread (**g**) and delay (**h**) of first spike vs recorded layer in sparse (orange) and dense (blue) networks. In **e.-f.** all data are shown and the mean ± SEM is represented as thick lines. Data in **c.-h.** are from n = 15 sparse networks of density = 347 ± 81 neurons/mm^2^ and from n = 16 dense networks of density = 686 ± 122 neurons/mm^2^.

There were two basic modes of propagation, depending on the density of the networks and stimulus parameters (Fig. 2a-d). In the temporal mode, most often observed in sparse networks (density ≲450 neurons/mm^2^) or with a small number of stimulated neurons, a narrow stimulus packet delivered to 20-40 ROIs in the 1^st^ layer evoked time-locked responses. Repeated delivery of the stimuli evoked action potentials that exhibited little trial-to-trial jitter as seen from the dot rasters and poststimulus time histograms (PSTSHs) (Fig. 2a,c). In subsequent layers, however, the firing became less reliable: the firing rate (average in a 500 ms window after stimuli) and probability (of at least 1 spike) decreased to nearly 0 by the 4^th^ layer (Fig. 2e,f, orange). The jitter in spike times remained small and was manifested as a slight widening of the spike packet (≲5 ms) in successive layers (Fig. 2g, orange). Hence, temporal information was preserved but only for 2-3 layers (see also below).

Propagation in the temporal mode was primarily feedforward despite the presence of reciprocal connections between layers. The spike onset increased linearly up to the 3^rd^ layer with a slope of 10.5 ms/layer (Fig 2h). Three factors contributed to this delay between layers: 1) a delay of 5 ms as the action potential propagated across the 1.1 mm interlayer gap (conduction velocity of 200 μm/ms in these experimental conditions (Patolsky, Timko et al. 2006, Barral and Reyes 2016)); 2) a synaptic delay of 3 ms (Muller, Swandulla et al. 1997, Barral and Reyes 2016); and 3) a delay of few ms due to the rise in membrane potential towards threshold (membrane time constant ~20 ms for neurons in culture (Barral and Reyes 2016)). The sum of these 3 factors is comparable to the delay measured experimentally. Hence, the propagation across layers was mediated by monosynaptic feedforward connections, with minimal contribution from recurrent and feedback connections. Simple feedforward models (Diesmann, Gewaltig et al. 1999) generally capture the dynamics of the temporal propagation mode with the exception that there was not a stable propagation regime (attractor) in our biological network. Increasing the number (spike packet amplitude) and/or decreasing the jitter (packet width) did allow propagation across all layers albeit asynchronously, in stark contrast to model predictions (see below).

The second mode of propagation, the rate mode, was most prominent in dense networks (≳450 neurons/mm^2^) or when a large number of neurons were stimulated synchronously and differed from the temporal mode in several regards. In response to a single packet in the first layer, firing in the stimulated layer had a transient and a persistent component (Fig. 2b,d). The transient component resembled the response in sparse networks above: the spikes had little trial-to-trial jitter and the PSTH matched the time course of the pulse packet. After a short silent period, the persistent component appeared where neuron fired multiple action potentials that lasted several hundred ms (Fig. 2b,d). Unlike the transient component, the action potentials in the persistent phase exhibited substantial trial-to-trial variability. Whereas the transient component completely disappeared by the 3^rd^ layer, the persistent component propagated reliably to layer 4 (Fig. 2b,d). Both the firing rate and probability remained high across layers (Fig. 2e, blue). Plotting the firing rate averaged over all layers (Fig. 2f) or spike probability in layer 4 (inset) vs density shows that this mode of propagation occurred reliably in networks with densities ≳450 neurons/mm^2^. In dense networks, the delay in the first action potentials increased with layer at a high rate (35.9 ms/layer) (Fig. 2h), indicating that propagation involved polysynaptic, recurrent connections within and between layers (see below). Though the persistent component propagated, temporal information was lost as jitter (or spread) of the first evoked spike taken over several trials increased substantially with layer (Fig. 2g).

### Propagation of temporal information

To further study how temporal information was propagated, the pulses were temporally jittered so that the summed activity was Gaussian distributed (so-called ‘pulse packets’ (Aertsen, Diesmann et al. 1996, Diesmann, Gewaltig et al. 1999)). The pulse width (jitter) was systematically varied (Fig. 3a and Supplementary Fig. 3) to evoke high probability firing with different spreads in layer 1 and to determine the optimal conditions for propagation of activity. Synchronized pulse packets resulted in higher firing rates and increasing the jitter caused a decrease in firing rate by about 50% and 30% in sparse and dense networks, respectively (Fig. 3b-e).

Tracking the evolution of firing probability (of at least 1 spike) and spread in successive layers demonstrated the tradeoff between propagation reliability and temporal fidelity (Fig. 3f). In the rate mode (blue), the trajectories converged in layer 2 regardless of initial values and the spread continued to increase with successive layers. It is noteworthy that the trajectories differed from theoretical predictions. Networks undergoing rate mode propagation did not display stable non-zero attractors with low jitter and high spiking probability. Rather, the attractor occurs at a very large spike packet width that extends beyond the conditions of the pulse packet (Fig. 3), which suggest a different mechanism than the one predicted by theory (Diesmann, Gewaltig et al. 1999, Reyes 2003).

In contrast, the trajectories for the temporal mode (orange) were nearly perpendicular: although the probability decreased, the increase in jitter was relatively small. In the temporal mode, the trajectories remained separated (Diesmann, Gewaltig et al. 1999), indicating that differences in the spread in deeper layers more reliably reflected the jitter in the first layer. Interestingly, differences between rate and temporal mode not only depended on the network architecture but also on the input: uniformly and synchronously stimulating all neurons within the field of view (0.7 × 1.5 mm^2^) allowed propagation for both sparse and dense networks (red and dark blue lines in Fig. 3f, respectively).

**Figure 3:**
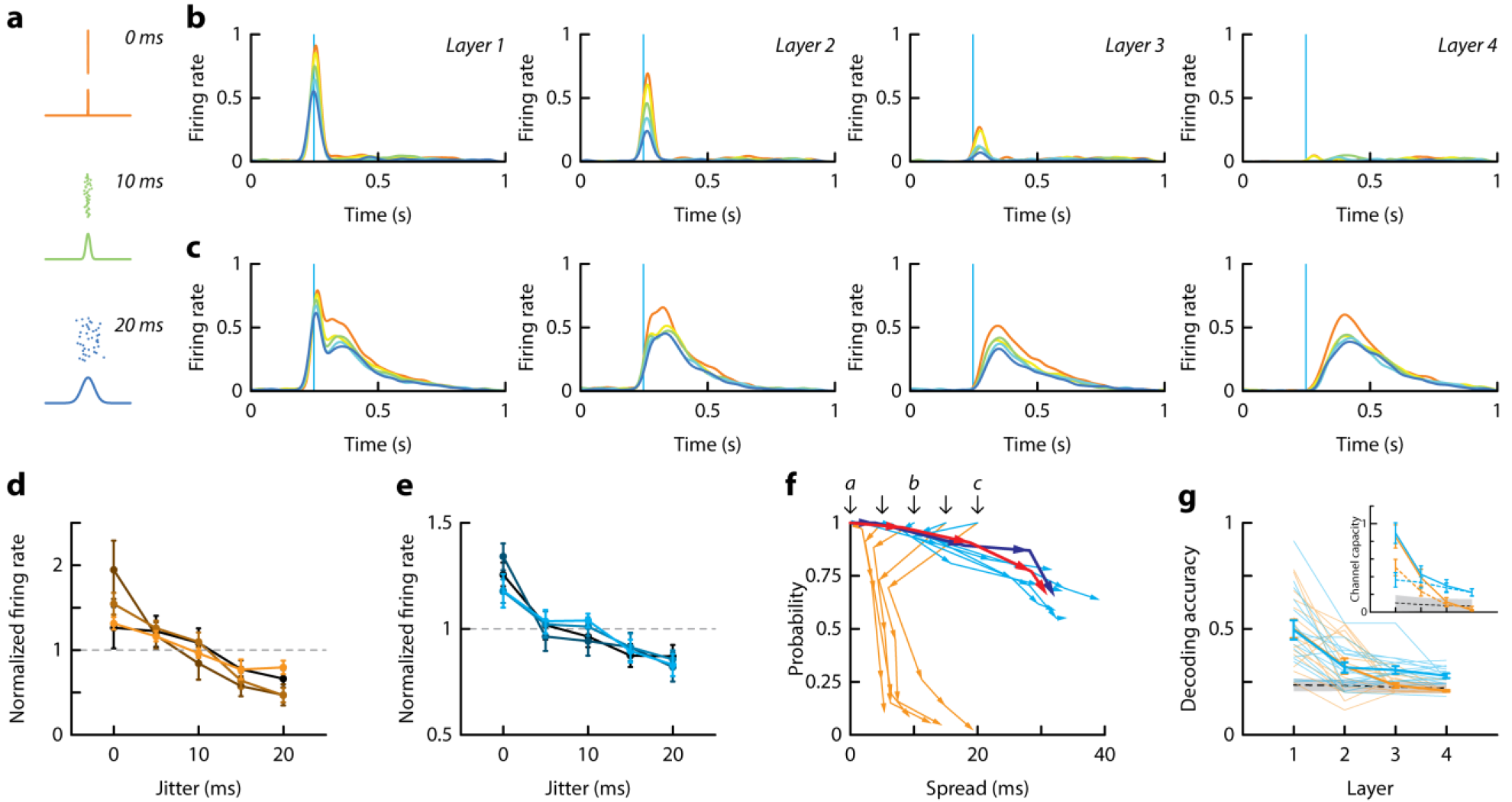
Propagation of spike timing in the feedforward networks. **a.** Examples of initial pulse packets of different widths. **b.-c.** Average firing rate for different stimulus jitters (red to blue for 0 to 20 ms jitters) in sparse (**b**) and dense (**c**) networks. **d.** Normalized firing rate as a function of jitter for neurons in layer 1 to 4 (from orange to black, respectively) of sparse networks. For each neuron in each network, the firing rate in the 150 ms following the stimulus was normalized to its mean for different jitters. **e.** Same as in **d.** for dense networks. **f.** Phase diagram of spike propagation. Spike probability is plotted as a function of spike spread for different initial conditions (indicated by black arrows: initial pulse packet of 0 to 20 ms width) in sparse (orange) and in dense (cyan) networks. Black arrows indicate values in layer 1 and arrowheads connected by lines indicate values in successive layers. Red and dark blue lines denote trajectories in sparse and dense networks, respectively, where all neurons in the field of view were activated. Data in orange and cyan are from the same networks as in Fig. 2. Red and dark blue trajectories are from different experiments and are from n = 17 sparse networks of density = 303 ± 59 neurons/mm^2^ and from n = 14 dense networks of density = 569 ± 80 neurons/mm^2^, respectively. **g.** Decoding accuracy measured as the probability of attributing 2 features of neuronal activity (time of the first spike and firing rate) to the correct stimulus vs layer for sparse (orange) and dense (blue) networks. Inset: channel capacity computed using the time of the first spike and the firing rate (solid lines) or the firing rate only (dotted lines) as features to the decoder. Data are from the same networks as in Fig. 2 and are presented as mean ± SEM. In **g.** all data are shown as thin lines.

To determine whether neuronal firing contained information about the stimulus, we trained a classifier to identify the stimulus (see Methods). We used both the time of the first spike and the firing rate in a 150 ms window after the light stimulation to identify the initial jitter. In the temporal mode, decoding accuracy was higher in the first layer but dropped to chance level by the 3^rd^ layer whereas it remained above chance in the rate mode (Fig. 3g). To directly quantify how much information could be propagated, we also computed the channel capacity (inset in Fig. 3g). In the first layer, we measured a channel capacity of ~0.8 bit in sparse or dense networks which represents about one third of the upper bound defined as the total entropy of the stimulus (~2.3 bits). In subsequent layers, this measure decreased to less than 0.1 bits in layer 4 of sparse networks but remained above 0.2 bits in dense networks. Decoding using firing rate as the unique feature resulted in degraded performance in the 1^st^ and 2^nd^ layers but decoding was almost identical for the 3^rd^ and 4^th^ layer suggesting that temporal information was propagated as firing rate in the rate mode in the deeper layers.

### Propagation of rate information

To examine whether different firing frequencies in the first layer could be discriminated in the deeper layers, data from dense network only were combined across different jitter stimuli and sorted according to the firing rate in the 1^st^ layer (in a 0.5 s window after the light stimulus). To compare quantitatively this analysis with the previous one, we generated 5 groups of equal number of trials. Plotting the firing rate for these 5 groups in the different layers showed that the separation artificially generated in layer 1 was faithfully propagated up to the 4^th^ layer (Fig. 4a), suggesting that firing rate was propagated in the rate mode.

To further characterize rate propagation, we constructed logistic maps where the abscissa was firing rate in a given (*n*th) layer and the ordinate is the firing rate in the next (*n*+1th) layer (Fig. 4b). For frequencies ≲15 Hz, the curve (averaged from 16 networks) superimposed with the unitary slope line, indicating that in this range, firing rate was preserved across layers and uniquely represented. At higher frequencies, the curve became sublinear, which indicates that firing rate decreased in successive layer to a fixed value of ~15Hz (see trajectory in Fig. 4b). We then grouped and averaged the data based on the firing rate in the first layer and traced the firing rate in successive layers (Fig. 4c). The traces remained separated in layer 4, indicating that the frequencies can be discriminated. Finally, we delivered 0.5 s-long Poisson trains of light pulses to the first layer with various input frequencies and observed that the firing rate in the 4^th^ layer increased linearly with that of the input delivered to the first layer (Supplementary Fig. 4).

**Fig. 4:**
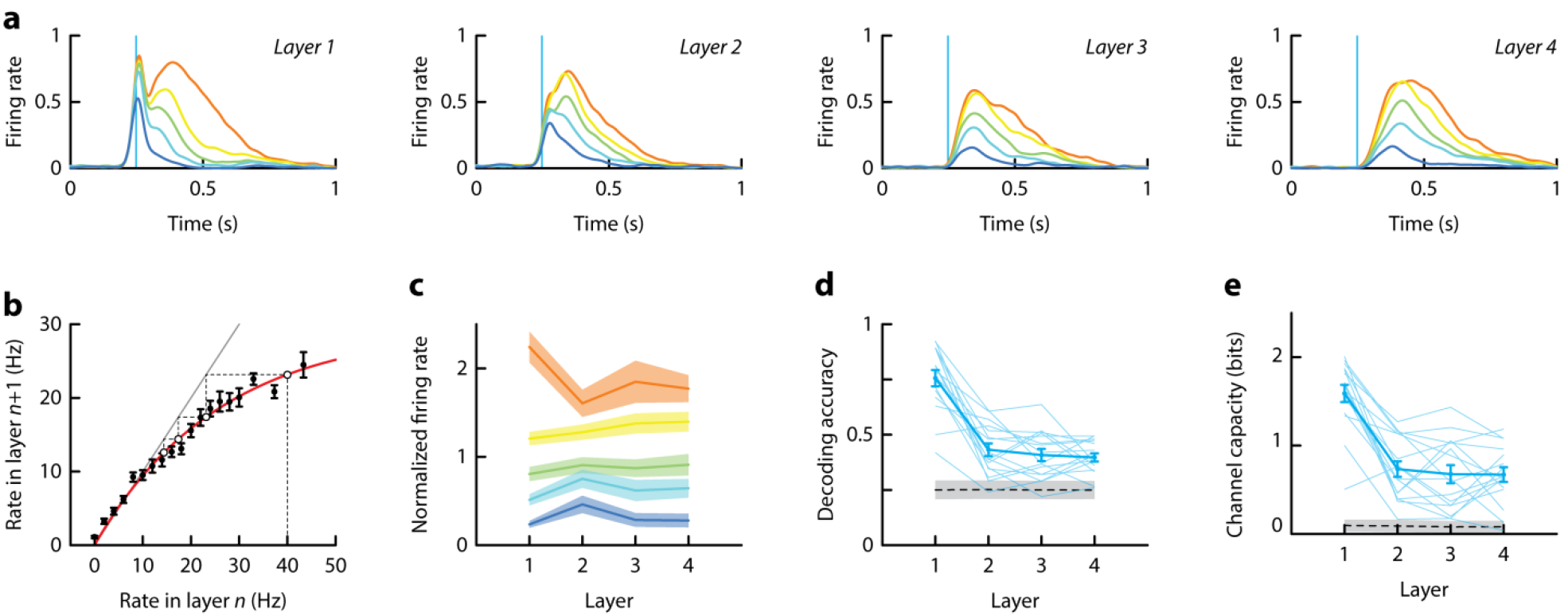
Propagation of firing rate information in dense networks. **a.** Firing rate in dense networks. For each network trials were sorted in 5 groups according to firing rate in layer 1 and averaged. **b.** Logistic map showing output rate (= firing rate in layer *n*+1) versus input rate (= firing rate in layer *n*) in dense networks. The firing rate was the average firing rate during 500 ms after stimulus. The grey line denotes slope of unity and the red line is a fit to the data. Dotted lines represent a sample trajectory starting at 40 Hz in layer 1 and ending at 13 Hz in layer 4. **c.** Normalized firing rate vs recorded layer. Data were separated in 5 groups according to their firing rate in layer 1. **d.** Decoding accuracy measured as the probability of attributing a given firing rate measurement to the correct group vs layer. **e.** Channel capacity computed using the decoded firing rates. Data are from the same dense networks as in Fig. 2 and are presented as mean ± SEM. In **d.-e.** all data are shown as thin lines.

To quantify the extent to which firing rates in the 1^st^ layer can be discriminated in successive layers, we trained a classifier and defined the decoding accuracy as the probability of correctly identifying the firing rate in layer 1 based on the firing rate in the *n*th layer (firing rate in the 0.5 s following the stimulus, see Methods). Decoding accuracy was greatest in layer 1 and decreased in successive layers but remained well above chance (Fig. 4d). To directly quantify how much information could be propagated in our system, we computed the channel capacity (Fig. 4e). This measure peaked at a value of about 1.6 bits in layer 1, which correspond to the maximum capacity of information that a single neuron is able to encode. In layers 2-4, channel capacity was constant and attained a value of about 0.7 bits.

For an effective rate code, the correlations in the firing of neurons should be low (Zohary, Shadlen et al. 1994). To measure correlations, we performed cell-attached recordings from pairs of neurons within and between layers. To measure noise correlation between neurons, we subtracted the ‘signal’ correlograms, constructed from shuffled trials, from the raw correlograms (see Supplementary Figs. 5 and 6). The noise correlation measured at the peak was low for neurons in the same layer (median value of C = 0.03; 1^st^ and 3^rd^ quartile [-0.01, 0.17]) and for neurons separated by 1, 2, or 3 layers (median values of C = 0.02; 1^st^ and 3^rd^ quartile [-0.05, 0.08]).

### Maintaining *E-I* balance during propagation

To examine the synaptic potentials underlying activity propagation, we performed whole-cell recordings from neurons in layer 1-4 and stimulated the 1^st^ layer (Fig. 5a,b). To maximize propagation, the light pulses were delivered synchronously to all pyramidal neurons within the field of view (0.7 × 1.5 mm^2^). In low-density networks where the evoked firing lacked a persistent component and did not propagate reliably (cf Fig. 2a,c), the postsynaptic potentials (PSPs) recorded in the first layer (Fig. 5a, black) exhibited a sharp rise to peak and then a slow decay back to baseline. In the next layers, the amplitude and decay rate decreased while the delay and rise time increased.

In networks in the rate propagation regime where the firing had prominent persistent component (cf Fig. 2b,d), the underlying PSPs in the first layer rose rapidly to a peak but decayed at a slower rate compared to the non-propagating network (Fig. 5b). There was a large decrease in PSP amplitude in the second layer followed by smaller decreases in subsequent layers. The evoked firing mirrored these changes in PSPs (Fig. 2b,d vs Fig. 5a,b). In the first layer, the sharp rise in the PSP amplitude accounted for the transient firing while the slow decay produced the persistent component. The delay between the two components is likely due to synchronized recurrent inhibition in the first layer. In the following layers, the peak of the PSTHs lagged the peak of the PSP because the smaller amplitudes required a longer integration time for the membrane potential to cross threshold.

The balanced regime –where the E and I synaptic inputs track each other both in magnitude and in time (Okun and Lampl 2008, Renart, de la Rocha et al. 2010, Barral and Reyes 2016)– is maintained during propagation within and across layers. To view the E and I synaptic potentials, we held the membrane potentials at -80 or 0 mV (reversal potentials of I and E, respectively) during repeated delivery of identical stimuli (Fig. 5c). In the first layer, the large, sharp peak in the composite PSP measured at resting potential (Fig. 5c, middle, black) was due to a combination of a rapidly rising EPSP (bottom, black) and an IPSP (top, black). The IPSP increased nearly proportionately with the EPSP (Fig. 5d,e) but with a delay (Fig. 5f). The IPSP increased the decay rate of the composite PSP but did not cancel the depolarization (Fig. 5c, middle). In subsequent layers, the EPSP and IPSP amplitudes both decreased (Fig. 5d,e); though the differences were small, the EPSPs decreased at a faster rate than the IPSPs. The magnitude of the E and I potentials at each layer co-varied (Fig 5d), consistent with the balanced regime.

**Fig. 5:**
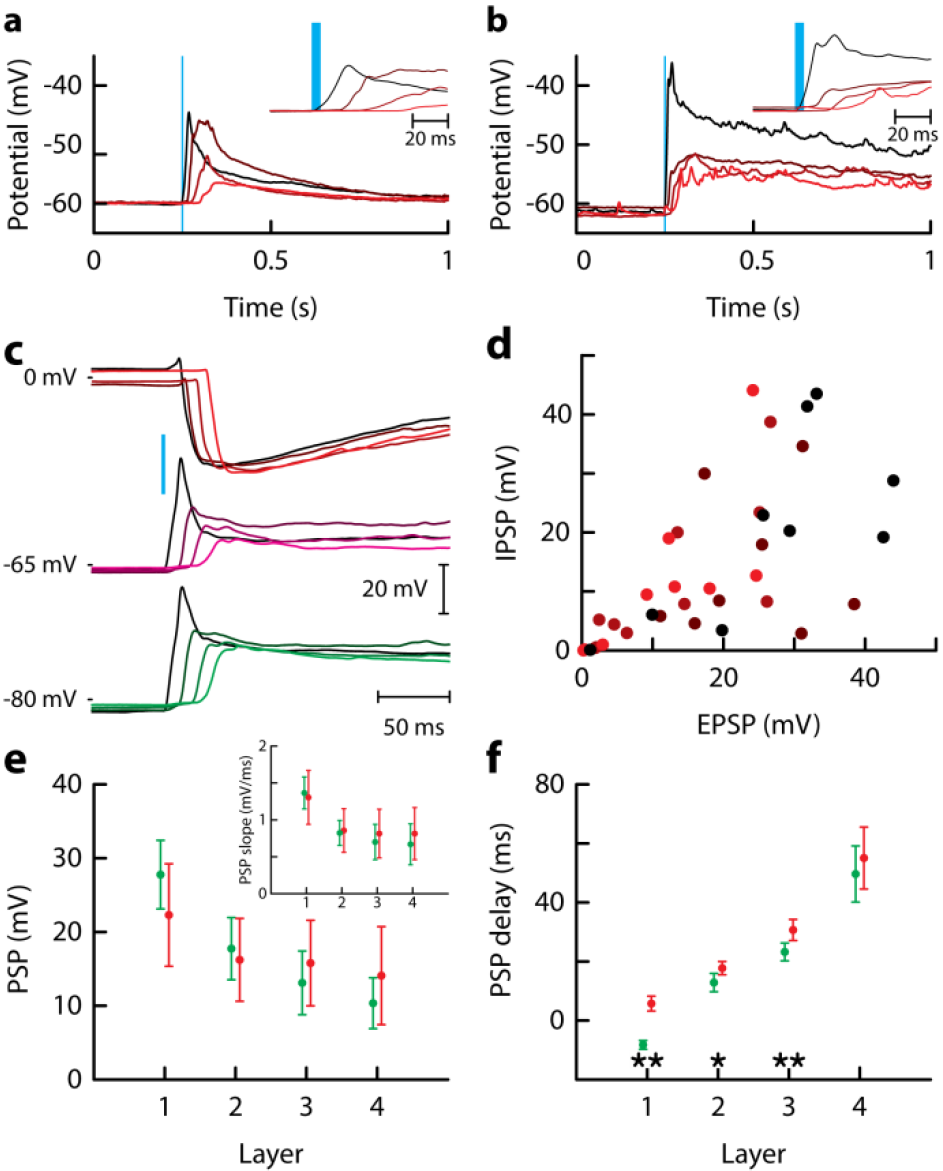
PSPs in feedforward networks. **a.** Average postsynaptic potential of neurons in sparse networks (n = 6 networks, density = 342 ± 57 neurons/mm^2^, mean ± STD). In each network, a single neuron was recorded in each layer (from black to red for layer 1 to 4, respectively; same color code in **a.-b.**). **b.** Same as in **a** but for dense networks (n = 5 networks, density = 572 ± 117 neurons/mm^2^, mean ± STD). **c.** Example of recordings of excitatory (EPSP, green), inhibitory (IPSP, red), and compound post-synaptic potentials when the same neuron was recorded in response to stimulation of different layers. **d.** IPSP vs EPSP for different layers (from black to red for layer 1 to 4, respectively; n = 9 networks where a neuron was recorded in each compartment, density = 444 ± 146 neurons/mm^2^, mean ± STD). Pearson correlation between EPSPs and IPSPs was 0.64 (*p* = 2.3×0^−5^, *n* = 36). **e.** PSP size vs layer (n = 6 networks where a neuron was recorded in each compartment, density = 381 ± 53 neurons/mm^2^, mean ± STD). Inset: maximal slopes of PSP. **f.** PSP delay measured as the time of maximal slope. Data in **e.** and **f.** presented as mean ± SEM.

The relative timing of the EPSPs and IPSPs was also preserved across layers. We measured the delays of EPSPs and IPSPs at the maximal slope of the membrane potential. In the first layer, there was a relatively long delay between the EPSPs and IPSPs because only the E cells were stimulated and some time was needed for the inhibitory cells to fire (Fig. 5f). The result is that there is a time window where the edge of the EPSP can ‘escape’ inhibition to evoke the early spikes of the transient component (Fig. 2a-d). In layers 2-4, the EPSP-IPSP delay was significantly shorter as both E and I cells could receive afferents from both E and I cells in the previous layer (Fig. 5f). With no sharp peaks in the composite PSP, the evoked action potentials were delayed (Fig. 2b,h). Thus the amplitude, shape and arrival time of EPSPs were well matched by those of IPSPs, producing a tight balance between excitation and inhibition that impeded further propagation of synchronous input in deeper layers.

We hypothesized that the prolonged activity in the rate propagation mode was due to a combination of recurrent activity and an NMDA synaptic current. Using paired recordings, we confirmed that a strong NMDA-mediated component in the E synapses between pyramidal cells in cultures (Supplementary Fig. 7). When the NMDA currents were blocked with APV, the persistent component disappeared and only the transient component propagated albeit incompletely (Fig. 7a top vs middle, see also Supplementary Fig. 8). Propagation was restored when the inhibition was also blocked with bicuculline as shown by the increase of spike probability and firing rate in all layers (Fig. 7a bottom; Fig. 7b,c). However, although several spikes could be observed in all layers, the persistent activity was shorter and could not be fully recovered (Fig. 7a bottom, see also Supplementary Fig. 8).

**Figure 6:**
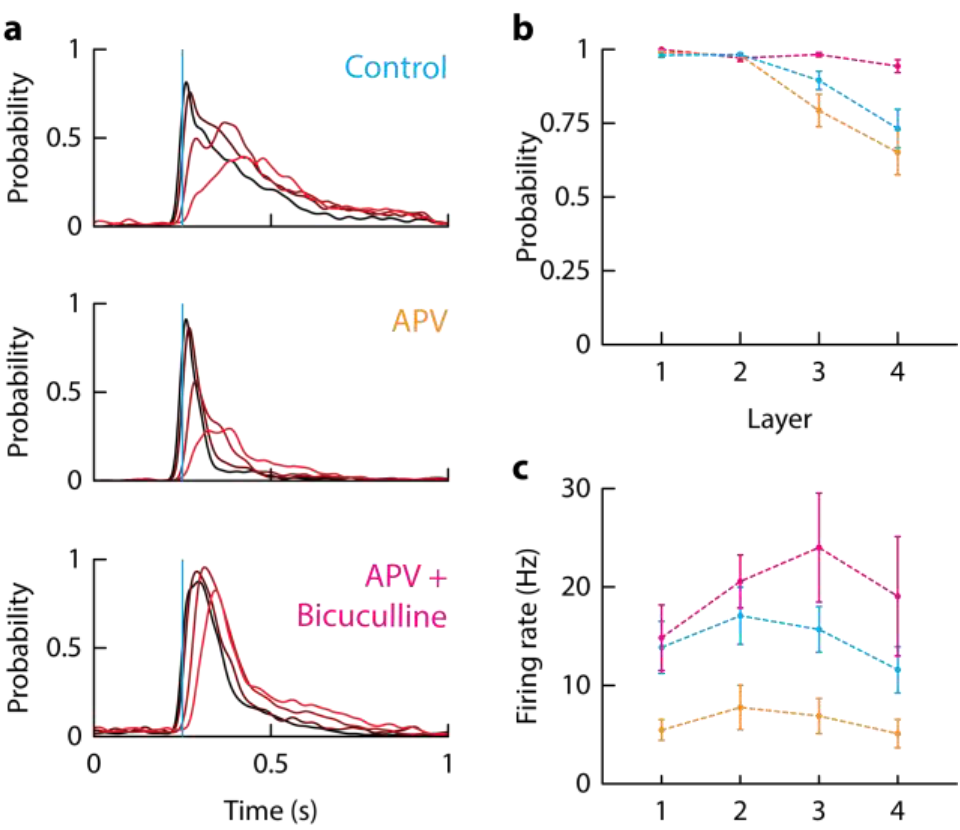
Role of AMPA and NMDA synapses in the feedforward propagation. **a.** Peristimulus time histogram showing firing rate recorded from a neuron in each layer (layer 1 to 4 from black to red; n = 9 networks, density = 278 ± 68 neurons/ mm^2^ - mean ± STD). Results shown for control condition (top), with the addition of 50 μM APV (middle) and with the addition of 50 μM APV and 10 μM bicuculline (bottom). **b.** Spike probability as a function of layer for the different conditions. **c.** Firing rate as a function of layer. Data presented as mean ± SEM.

### Propagation in heterogeneous networks

The fact that propagation occurs more readily in networks where recurrent connections were sufficiently dense to support persistent activity, suggests that the direction of propagation may be biased in the direction of increasing network size. In cortex, for example, the number of neurons in progressively higher order regions varies significantly (Schuz and Palm 1989, Collins, Airey et al. 2010). We therefore constructed a feedforward chamber where the area of each compartment increased in successive layer (Fig. 7a). Using the spatial profile of connection probability (Supplementary Fig. 1), we estimated that the number of connections *K* increases by more than 3 fold from 1^st^ to 4^th^ layer (Supplementary Fig. 2). Neurons in the small layer were stimulated synchronously and the activity of neurons monitored in the direction of increasing layer size (Fig. 7b); then, an equal number of neurons were stimulated in the large end and spiking activity monitored in the direction of decreasing layer size (Fig. 7c).

**Figure 7:**
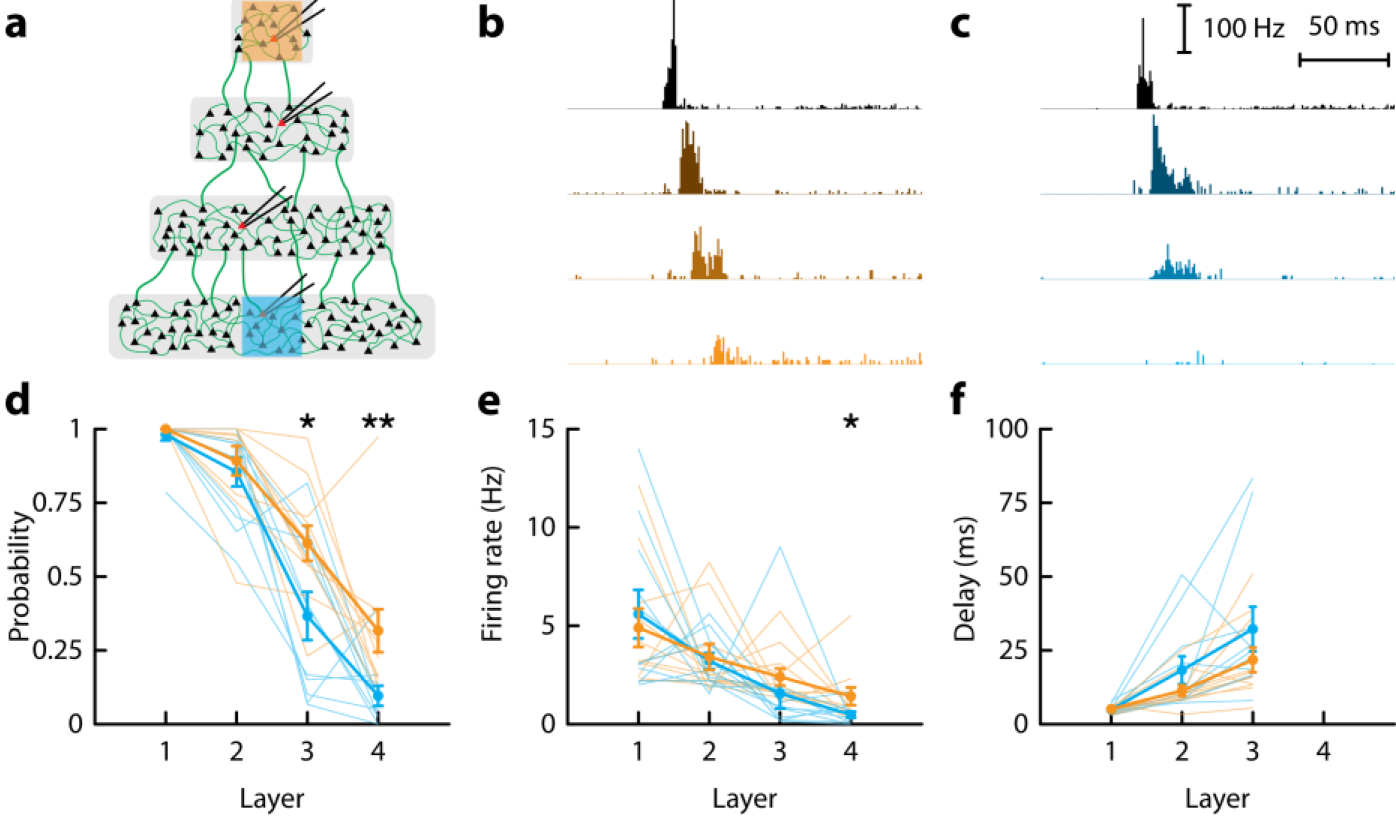
Propagation in networks of different sizes. **a.** Schematic of the feedforward network with layers of different sizes (n = 11 networks, density = 297 ± 111 neurons/mm^2^, mean ± STD). An area of same dimension was photostimulated either in the smallest or in the largest network as depicted. **b.-c.** Peristimulus time histogram of neurons when the small (**b**) or the large (**c**) network was stimulated. In each network, a single neuron is recorded in each layer (from black to blue (orange) for layer 1 to 4, respectively). **d.-f.** Spike probability (**d**), firing rate (**e**), and delay of first spike (**f**) as a function of layer when the smallest (orange) or the largest (blue) network was activated. In **d.-f.** all data are shown and the mean ± SEM is represented as thick lines. Paired *t*-test statistics \**P* < 0.05, \*\**P* < 0.01.

As anticipated, propagation occurred more readily in the direction of increasing layer size. This was shown by increased spike probability and firing rate in the last layer (Fig. 7d-e). The delays of action potentials were slightly shorter in the direction of increasing size (Fig. 7f), consistent with the fact that small networks resembled sparse networks. Taken together, these results suggest that the direction of signal transmission could be amplified and biased in non-homogeneous networks.

## Discussion

We found that the density of neurons in a biological model of feedforward network could determine the type of information that was propagated. Sparse networks were characterized by 1) a phase diagram that resembled the one of ideal feedforward networks with a low number of neurons in each layer (Diesmann, Gewaltig et al. 1999); 2) a short delay of propagation that was in agreement with a single synapse between layers and thus suggested that the network could be reduced to its feedforward connections only; and 3) by a temporal spikes precision that was low and constant which is a necessary condition for propagation of temporal information. In dense networks, although spikes were reliably propagated, temporal information was lost in contradiction with previous theoretical results (Diesmann, Gewaltig et al. 1999). The long delay suggested that these networks could be represented rather as the assembly of recurrent modules than as an ideal feedforward network. In this case, firing rate information was instead well transmitted from one layer to the next and this propagation did not result in the increase of synchrony, a prerequisite for propagating a firing rate code (Zohary, Shadlen et al. 1994). A decorrelation mechanism must therefore be in place. Our results showed that excitation and inhibition were balanced suggesting that mechanisms based on tracking and cancellation of excitatory and inhibitory inputs could take place (Renart, de la Rocha et al. 2010). It is also possible that heterogeneity in intrinsic (Padmanabhan and Urban 2010) and synaptic properties, recurrent activity, or a combination of both mechanisms permit to achieve asynchronous propagation.

Simulations have so far been the only way to study feedforward networks in a systematic manner. Several refinements of the purely feedforward network have been developed to better account for the architecture of biological networks. For example, feedback connections were introduced between layers and were shown to stabilize the propagation of synchronous inputs (Moldakarimov, Bazhenov et al. 2015). To model networks in more natural settings, the feedforward chain was embedded in a larger recurrent network (Aviel, Mehring et al. 2003, Mehring, Hehl et al. 2003, Tetzlaff, Buschermohle et al. 2003, Vogels and Abbott 2005, Kumar, Rotter et al. 2008, Jahnke, Memmesheimer et al. 2013, Chenkov, Sprekeler et al. 2017). Several lines of experiments support the model of feedforward networks in biological systems. Propagation of firing rate has been demonstrated in the drosophila olfactory system (Jeanne and Wilson 2015). Propagation of temporal information has also been studied *in vivo* in the auditory system of the locust (Vogel and Ronacher 2007) or in the song generation system of the zebra finch bird (Kimpo, Theunissen et al. 2003, Long, Jin et al. 2010). However the actual input was never known in these experiments. Here, we provide the first experimental study of a feedforward network where the input delivered to the network could be controlled faithfully and varied systematically. Interestingly, we found two modes of propagation: a purely feedforward mode and another mode that requires recurrent activity.

Many parameters vary from lower to higher cortical areas whether it is the cell density (Schuz and Palm 1989, Collins, Airey et al. 2010), intrinsic properties of neurons (Gilman, Medalla et al. 2016) or connectivity (Elston 2000, DeNardo, Berns et al. 2015). Here, we reproduced a hierarchical arrangement of cortical modules by building a feedforward network with a gradient in the number of neurons. The linear increase in size is ultimately linked to a sublinear decrease in the synaptic strength (Barral and Reyes 2016) such that the total input increases in the direction of larger networks. This asymmetry may lead to signal amplification and provides a direction of better propagation that could mimic the relation between bottom-up and top-down signal propagation.

We previously showed that synaptic strengths scale with network size to preserve balance between excitation and inhibition, maintain variable spiking statistics and reduce correlations in spiking as predicted by theory and observed in the intact brain (Barral and Reyes 2016). Here, we provide a functional significance for synaptic scaling. Our results indicate that spike propagation depends quantitatively and qualitatively (timing vs rate) not only on the input pulse packet but also on network architecture and density. Thus synaptic scaling controls the neural code in play for the propagation of information in modular networks.

## Acknowledgement

We thank M. Weck for the use of the clean room and T. Pinon and J. Palacci for help with the microfabrication process. This work was supported by a NIH grant R01DC005787 (AR). XJW was supported by Naval Research grant N00014-17-1-2041, NIH grant R01MH062349, Science and Technology Commission of Shanghai Municipality grant 15JC1400104. JB was supported by a Human Frontier Science Program long-term postdoctoral fellowship (LT000132/2012) and by the Bettencourt Schueller Foundation.

## Author Contributions

JB, XJW and AR designed the project. JB performed the experiments and analysed the results. JB, XJW and AR wrote the manuscript.

## Materials and Methods

### Primary neuron cultures

Dissociated cortical neurons from postnatal (P0-P1) mice were prepared as described previously (Hilgenberg and Smith 2007, Barral and Reyes 2016) and in accordance with guidelines of the New York University Animal Welfare Committee. Briefly, the mouse cortex was dissected in cold CMF-HBSS (Ca^2+^ and Mg^2+^ free Hank’s balanced salt solution containing 1 mM pyruvate,15 mM HEPES, 10 mM NaHCO_3_). The tissue was dissociated in papain (15 U/mL, Roche) containing 1 mM L-cystein, 5 mM 2-amino-5-phosphonopentanoic acid and 100 U/ml DNase (DN25; Sigma) for 25 min. After enzymatic inactivation in CMF-HBSS containing 100 mg/mL BSA (A9418; Sigma) and 40 mg/mL trypsin inhibitor (T9253; Sigma), pieces were mechanically dissociated with a pipette. Cells concentration was measured before plating using a haemocytometer. Approximately 0.3-3×10^6^ cells were plated on each coverslip, resulting in a density of ~100-1,000 cells/mm^2^ at the time of experiment. Neurons were seeded onto German glass coverslips (25 mm, #1 thickness, Electron Microscopy Science). Glass was cleaned in 3 N HCl for 48 h and immersed in sterile aqueous solution of 0.1 mg/mL poly-L-lysine (MW: 70,000 – 150,000; Sigma) in 0.1 M borate buffer for 12 h. Neurons were grown in Neurobasal medium (supplemented with B27, Glutamax and penicillin/streptomycin cocktail; Invitrogen) in a humidified incubator at 37 °C, 5% CO_2_. One third of the culture medium was exchanged every 3 days.

Expression of channelrhodopsin (ChR2) in excitatory neurons was achieved by crossing homozygote *Vglut2-Cre* mice (016963, Jackson Laboratory) with *ChR2-loxP* mice (Ai32, 012569, Jackson Laboratory). Experiments were performed at 14-21 DIV, when neuronal characteristics and network connectivity were stable and expression of ChR2 was sufficient to enable reliable photostimulation.

### Microfabrication and chip production

The micro-chambers comprise neuronal layers (width *w*, length *l*) for cell development that were separated by a gap of length *c* for axon growth. We designed a symmetric network where all layers had the same size (Fig. 1b; in mm: *w* = 0.7; *l* = 6; *c* = 0.4) and an asymmetric network where the length increased linearly with layer number (Fig. 7a; in mm: *w* = 0.5; *l* = 0.5, 1, 1.5, 2, *c* = 0.4) (see also Supplementary Fig. 2). The symmetric network was made of 6 layers and we recorded neurons in layers 2 to 5 (and not in layer 1 or 6) to avoid edge effects due to the fact that neurons from the terminal layers will make fewer connections (see Supplementary Fig. 2). For the asymmetric network, we built 7 layers and recorded from layer 1 to 4 to increase the variation of number of connections.

The chambers were made in PDMS using soft lithography and replica molding. To fabricate the master with positive relief patterns of cell culture compartments, we built a single layer of photoresist of 160 μm in height. A layer of SU82050 was spin-coated onto the wafer at 1200 rpm and then soft-baked for 7 min at 65°C and 30°min 95°C. The template was then exposed to UV light through an optic plastic mask (CAD/Art Services) of the culture compartment. After hard bake (5 min at 65°C, 12°min at 95°C, and 1 min at 65°C), the final mold was developed in SU8 developer.

We casted and cured a polydimethylsiloxane polymer (PDMS, Sylgard 184, Dow Corning) against the positive relief master to obtain a negative replica-molded piece PDMS was mixed with curing agent (10:1 ratio) and degassed under vacuum. The resulting preparation was poured onto the mold, pressed between two glass slides and cured at 110°C for 2 minutes onto a hot plate. After curing, the PDMS piece was peeled away, sterilized with ethanol and sealed onto the treated glass coverslip. The resulting assembly was washed with PBS and incubated at 37°C overnight. After rinsing, the device was flooded by culture medium. Neurons were added and cultured normally. After 24 hours, the PDMS mold was peeled away from the glass coverslip to allow processes to grow and connect different layers.

### Electrophysiological recordings

Recordings were performed at room temperature in HEPES-based artificial cerebrospinal fluid (aCSF). The aCSF solution contained (in mM): 125 NaCl, 10 NaHCO_3_, 25 D-glucose, 2.5 KCl, 2 CaCl_2_, 1.25 NaH_2_PO_4_, 1 MgCl_2_, and 10 HEPES. For some experiments, 50 μM APV was added to block the NMDA component of postsynaptic currents or 10 μM bicuculline to block inhibitory currents.

Electrodes, pulled from borosilicate pipettes (1.5 OD) on a Flaming/Brown micropipette puller (Sutter Instruments), had resistances in the range of 6-10 MΩ when filled with internal solution containing (in mM): 130 K-gluconate, 10 HEPES, 10 phosphocreatine, 5 KCl, 1 MgCl_2_, 4 ATP-Mg and 0.3mM GTP.

Cells were visualized through a ×10 water-immersion objective using infrared differential interference contrast (IR-DIC) and fluorescence microscopy (BX51, Olympus). Simultaneous whole-cell current-clamp recordings were made from up to four neurons using BVC-700A amplifiers (Dagan). The signal was filtered at 5 kHz and digitized at 25 kHz using an 18-bits interface card (PCI-6289, National Instrument). Signal generation and acquisition were here and in the following controlled by a custom user interface programmed with LabVIEW (National Instrument).

### Optical stimulation setup

We used a Digital Light Processing projector (DLP LightCrafter; Texas Instrument) to stimulate optically neurons expressing ChR2 as previously described (Barral and Reyes 2016, Barral and Reyes 2017). The projector had a resolution of 608×684 pixels. The image of the projector was de-magnified and collimated using a pair of achromatic doublet lenses (35 mm and 200 mm; Thorlabs). A dual port intermediate unit (U-DP, Olympus) was placed in-between the fluorescent port and the projection lens of the microscope and enclosed a 510 nm dichroic mirror (T510LPXRXT, Chroma). The resulting pixel size at the sample plane was a rectangle of dimensions 2.2 μm × 1.1 μm. We used the inbuilt blue LED of the projector which has a center wavelength of 460 nm and intensity of 10 mW/mm^2^ at the sample plane. The time resolution of the projector was 1,440 Hz.

### Stimulation and recordings protocols

We first selected *N*_*stim*_ = 15-60 regions of interest (ROIs) that were drawn onto ChR2 positive neurons (about 10-20 % of neurons that are in the field of stimulation). Care was taken to stimulate small areas of ~30 μm×30 μm to avoid stimulation of processes belonging to adjacent neurons. Each ROI was stimulated by a single light pulse of duration Δ_*pulse*_ = 5ms. The light intensity was fixed at 10 mW/mm^2^; a value that was sufficient to evoke reliably a spike in the selected neurons but too low to stimulate neighboring cells. Particular attention was paid to record neurons that do not express any ChR2 to avoid any obvious cross-activation.

In most experiments we focused on one experimental parameter: the dispersion crof the pulse packet. We used 5 values of temporal dispersion *σ* = (0, 5, 10, 15, 20 ms). In some experiments we used a.5 s-long Poisson trains of light pulses to the first layer and we varied the effective pulse rates (5, 10, and 20 Hz; Supplementary Fig. 4). A given stimulus was repeated *N*_*trias*_ = 5-6 times for PSPs data and *N*_*trias*_ = 10-40 times for spike data. We allowed at least 5 s of recovery between each stimulation. In each network, we recorded concurrently 4 neurons (1 neuron in each layer) in current clamp mode. Spike data were acquired in cell-attached configuration. When whole-cell configuration was established, we recorded the membrane potentials *V* in 3 different cases: (1) when *V* was close to the resting membrane potential, (2)-(3) when an injected current drove *V* close to the reversal potentials of excitation and inhibition, respectively.

## Data analysis

### Analysis of network characteristics

For every experiment, IR-DIC images around the region of recording were saved for off-line examination. Neuronal density *d* was determined by counting somata on a ~1 × 1 mm^2^ area in each layer. Network density was defined as the average density in each compartment. Data were pooled according to densities. Low and high densities corresponded to networks of neuronal densities (in neurons/mm^2^): *d* < 450, and 450 < *d*, respectively.

### Analysis of membrane potential data

Postsynaptic potential characteristics were computed after averaging PSPs over 5-6 repetitions. We determined the size of PSP at its maximum, the slope as the maximum of the PSP derivative, and the PSP delay as the time of maximal slope (see Fig. 5).

### Analysis of spike data

The spike probability was defined as the probability to observe at least 1 spike on a given trial (Fig. 2e-f). The delay and the spread were measured from the time of the first spike at each trial (Fig. 2g-h). The firing rate was defined as the average firing rate in a window of 150-500 ms following the stimulus, as mentioned in the main text (Figs. 2e-f, 3d-e, and 4b-c). Firing rates for peristimulus time histogram were computed by convolving the spikes data with a Gaussian kernel of width 25 ms (Figs. 3b and 4a).

### Decoding and mutual information

To decode temporal information (i.e. the correct stimulus among the different jitters *σ* ∈ Σ = [0, 5, 10, 15, 20 ms]), we trained a classifier for each neuron of each network to identify the stimulus using two parameters: the firing rate in the 150 ms following the stimulus and the time of the first spike (Fig. 3g). We used the Statistics and Machine Learning Toolbox from Matlab to fit the discriminant analysis classifier. This allowed us to estimate the conditional probability *p*(*r*_*l*_|*σ*) of observing the response *r* in layer *l* given the stimulus *σ*. We defined the decoding accuracy *d* as the average of the conditional probability over all stimuli (i.e. the proportion of well-attributed trials by the decode): *d_*l*_* = 〈*p*(*r*_*l*_|*σ*)〉_Σ_. This value was compared to the average value of 0.23 that was computed when trials were randomized. Note that this value is slightly higher than the lower bond 1/5 = 0.2 because the number of trials of each stimulus was not necessarily equal.

We used the non-uniform partitioning obtained from the classification results as the base to compute mutual information. Our method is equivalent to adaptive partitioning of the (*R*, Σ) space (Cellucci, Albano et al. 2005). Our partitioning maximizes decoding accuracy and thus the sum of diagonal elements of the matrix *p*(*r*_*l*_|*σ*) and limits bias in the estimation of mutual information. The mutual information *MI*_*l*_ between the stimulus and the firing rate in layer *l* was calculated as follow:

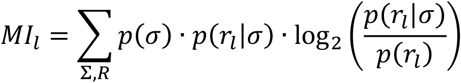

where *p*(*σ*) and *p*(*r*_*l*_) are the probability distributions of responses and stimuli, respectively, and *p*(*r*_*l*_|*σ*) is the stimulus conditional probability distributions of responses. In principle, the upper bound for mutual information is the entropy of the stimulus which equal log_2_(5) = 2.32 bits because the stimulus can take 5 different values. The channel capacity was then computed numerically using the Arimoto-Blahut algorithm (Cover and Thomas 2006). This method maximizes the mutual information and provides an estimate of how much information can be encoded.

In a second analysis, we used the firing rate as the sole feature for the decoder and calculated mutual information and channel capacity in the same way (inset in Fig. 3g).

To estimate how much information of the firing rate was propagated (Fig. 4), we ordered and grouped the firing rates measured in layer 1 into 5 non-overlapping groups and defined each group as a stimulus *s*. This was done for each network separately. We used the same number of stimuli (5) to be the same as the precedent analysis to compare quantitatively both results. Similarly, we the trained a classifier to discriminate and classify the stimuli *s* according to the firing rates in the layers 1-4. We measured the decoding accuracy and the channel capacity as described above.

### Statistical analysis

All the data were shown as mean ± SEM., unless stated otherwise. Two group comparisons were performed using either paired or unpaired two-sided Mann-Whitney *U*-test. Firing rates which had a long tail distribution were first log transformed before being compared. A generalized linear regression model with Bonferroni correction was performed when more than two groups were compared. The variances between groups were assumed to be different. No statistical methods were used to pre-determine sample sizes.

